# Bayesian inference of state feedback control parameters for *f*_*o*_ perturbation responses in cerebellar ataxia

**DOI:** 10.1101/2024.03.12.584554

**Authors:** Jessica L. Gaines, Kwang S. Kim, Ben Parrell, Vikram Ramanarayanan, Alvincé L. Pongos, Srikantan S. Nagarajan, John F. Houde

## Abstract

Behavioral speech tasks have been widely used to understand the mechanisms of speech motor control in healthy speakers as well as in various clinical populations. However, determining which neural functions differ between healthy speakers and clinical populations based on behavioral data alone is difficult because multiple mechanisms may lead to the same behavioral differences. For example, individuals with cerebellar ataxia (CA) produce abnormally large compensatory responses to pitch perturbations in their auditory feedback, compared to controls, but this pattern could have many explanations.

Here, computational modeling techniques were used to address this challenge. Bayesian inference was used to fit a state feedback control (SFC) model of voice fundamental frequency (*f*_*o*_) control to the behavioral pitch perturbation responses of individuals with CA and healthy controls. This fitting process resulted in estimates of posterior likelihood distributions of five model parameters (sensory feedback delays, absolute and relative levels of auditory and somatosensory feedback noise, and controller gain), which were compared between the two groups. Results suggest that the CA group may proportionally weight auditory and somatosensory feedback differently from the control group. Specifically, the CA group showed a greater relative sensitivity to auditory feedback than the control group. There were also large group differences in the controller gain parameter, suggesting increased motor output responses to target errors in the CA group. These modeling results generate hypotheses about how CA may affect the speech motor system, which could help guide future empirical investigations in CA. This study also demonstrates the overall proof-of-principle of using this Bayesian inference approach to understand behavioral speech data in terms of interpretable parameters of speech motor control models.

**Author summary:** Cerebellar ataxia is a condition characterized by a loss of coordination in the control of muscle movements, including those required for speech, due to damage in the cerebellar region of the brain. Behavioral speech experiments have been used to understand this disorder’s impact on speech motor control, but the results can be ambiguous to interpret. In this study, we fit a computational model of the neural speech motor control system to the speech data of individuals with cerebellar ataxia and that of healthy controls to determine what differences in model parameters best explain how the two groups differ in their control of vocal pitch. We found that group differences may be explained by increased sensitivity to *auditory feedback prediction errors* (differences between the actual sound speakers hear of their own speech as they produce it and the sound they expected to hear) and increased motor response in individuals with cerebellar ataxia. These computational results help us understand how cerebellar ataxia impacts speech motor control, and this general approach can also be applied to study other neurological speech disorders.

## Introduction

Altered auditory feedback experiments have been widely used to probe the mechanisms of speech motor control. In this class of experiments, participants speak while listening to a digitally-altered version of their own voice through headphones. This allows experimenters to observe how the speech motor system responds to a perceived error in some acoustic property of the voice such as pitch, formant frequencies, or loudness (for a review, see [1]). In a pitch perturbation task, the auditory feedback of participants’ production of a sustained vowel sound is, at an unexpected time and for a brief period (usually a fraction of a second), digitally altered to have a higher or lower fundamental frequency (*f*_*o*_; which is perceived as vocal pitch) than was actually produced.

Participants tend to compensate for this perceived error within the ongoing production by shifting their produced pitch in the opposite direction of the perturbation, demonstrating that people use auditory feedback for online control during speech.

Variations of this task have been used to inform the understanding of speech motor control in typical speakers [2–4] and, more recently, to compare the responses of healthy speakers with those of various clinical populations (e.g., Alzheimer’s disease [5]; Parkinson’s disease [6]; Hyperfunctional voice disorders [7]; Laryngeal dystonia [8]; Cerebellar ataxia [9, 10]).

However, due to the complexity of the speech motor system, it remains difficult to determine what specific mechanisms may lead to the observed differences in behavioral results in different populations. A difference in how two groups respond to a mid-trial pitch perturbation may create multiple hypotheses about the differences in the speech motor control system between the two groups. A mechanistic computational model can thus be a powerful tool in evaluating these hypotheses by simulating the effects of specific model changes on the observable output. Tuning the parameters of a computational model to fit observed behavioral data has the potential to distinguish how different model components contribute to the observed effect [11–16].

A recent example of mechanistic ambiguity in behavioral speech data is the increased response to pitch perturbations observed in individuals with cerebellar ataxia (CA; [9, 10]). CA is a condition characterized by a loss of coordination in limb movements, eye movements, and gait, as well as speech symptoms such as articulatory impreciseness, harsh or breathy voice, slowed speech, hypernasality, excessive variation of loudness, scanning speech, and/or impaired timing of voice onset [17, 18]. In CA these movement-related symptoms are caused by cerebellar lesions or degeneration, which is consistent with the idea of the cerebellum as a likely neural substrate for *internal models*, or neural representations of the body used in neuromotor control [19, 20]. However, open questions remain about the precise impact of cerebellar lesions on internal models and the speech motor control system.

Previous pitch feedback perturbation studies involving individuals with CA have examined the effects of CA on speech motor control. Houde et al. [9] found that individuals with CA displayed a heightened response to a 400 ms, mid-utterance pitch perturbation of 100 cents, with the peak of the CA group average response observed to be twice as high as that of the control group average. Around 300 ms after the end of the perturbation, however, the response of the CA group had fallen such that there was no significant difference between the average normalized pitch of the two groups during the time period from 0.7 to 1.0 s. The latency of the peak response was approximately the same for the two groups. These results are reproduced in Fig 1. Similar results were observed by Li et al. [10] using a slightly different paradigm with a 200 ms perturbation of 200 or 500 cents.

**Fig 1.**
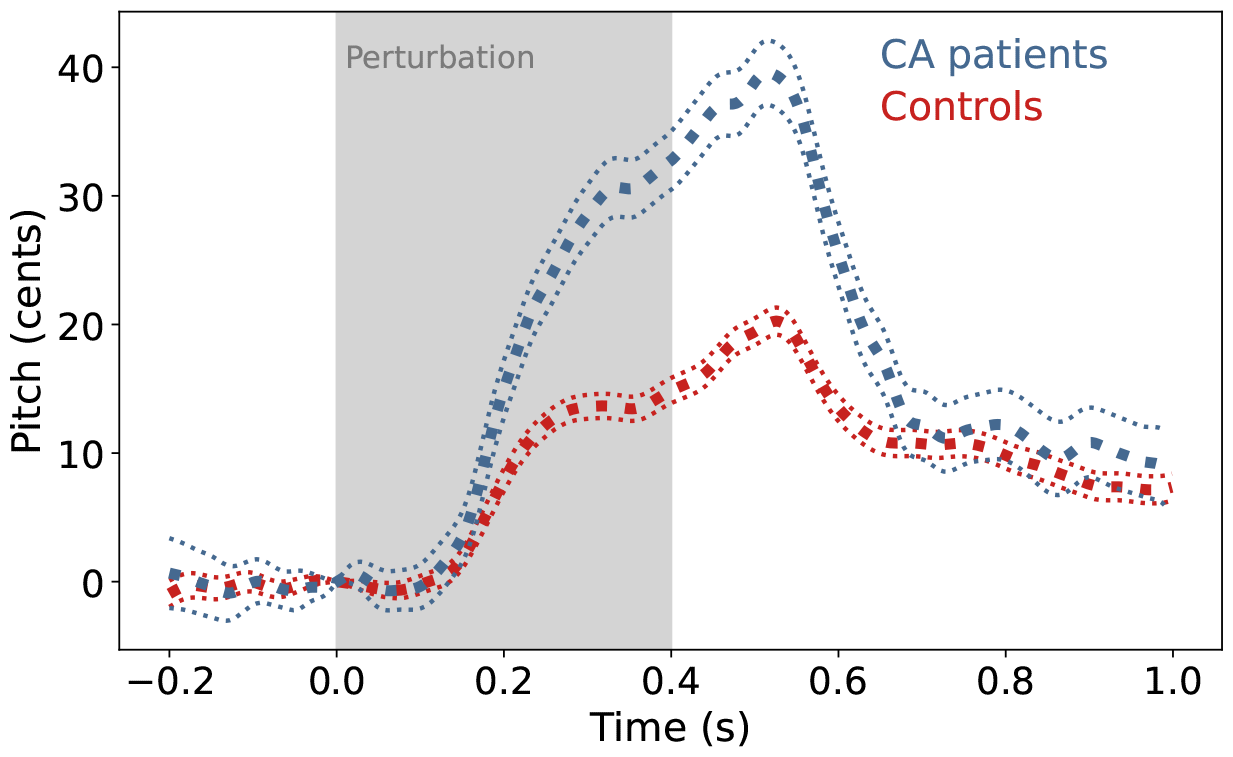
Behavioral data. Group averaged response (thick dotted line) to a 400 ms pitch perturbation of 100 cents for the CA group (blue) and control group (red) [9]. The thin dotted line indicates standard error. The CA group response showed a significantly larger magnitude than that of the control group with no change in peak latency.

Houde et al. [9] proposed two possibilities to explain these findings:

*Hypothesis 1* : Compared to the control group, the CA group exhibits increased reliance on auditory feedback in comparison with somatosensory feedback.

*Hypothesis 2* : Compared to the control group, the CA group displays an increased reliance on all types of sensory feedback collectively due to impairment of the feedforward system.

The current study further investigated these two hypotheses by estimating parameter values to fit a computational model of voice *f*_*o*_ to the empirical pitch perturbation responses of CA and control groups (as observed in Houde et al. [9]). A recently developed Bayesian inference method called simulation-based inference (SBI; https://sbi-dev.github.io/sbi/; [21, 22] was used to estimate posterior likelihood distributions for each model parameter based on each data set. The above hypotheses were investigated by comparing posterior probability distributions of parameters between the two groups and observing the impact of parameter ablation on the quality of model fit.

## Model

### Overview of state feedback control

The computational model used in this investigation implements a state feedback control (SFC) architecture to simulate a participant’s response to an unexpected pitch perturbation during a sustained vowel production. The theory of state feedback control has been well established as a plausible neural mechanism for non-speech motor tasks [23, 24] and has more recently been explored in the field of speech motor control [13, 25, 26].

This SFC model of voice control we consider is greatly simplified from the full task of voice control. Our model simulates only the control of *f*_*o*_ in ongoing voice output, which we idealize as controlling the rest-length of a spring in a single damped spring-mass system that is somewhat analogous to the cricothyroid muscle’s control of pitch [25, 27]. The controls generated by laryngeal motor cortex are modeled as desired changes in the rest-length of a single muscle determining *f*_*o*_ in our simplified larynx. The SFC architecture for motor control of this simplified larynx is shown in Fig 2.

**Fig 2.**
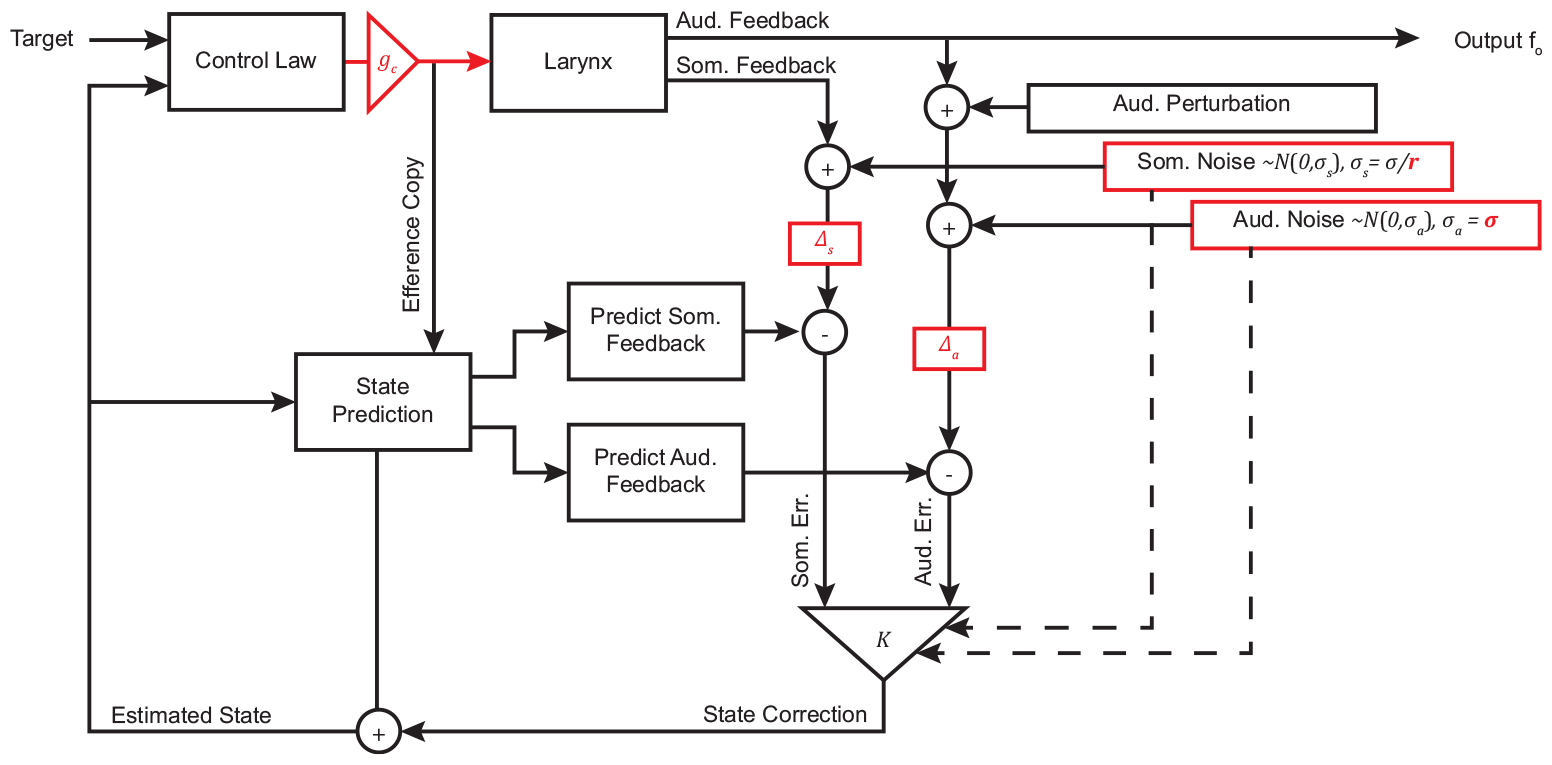
Overview of state feedback control. A state feedback model of vocal *f*_*o*_ control where Δ_*a*_ is auditory feedback delay, Δ_*s*_ is somatosensory feedback delay, *σ* is overall feedback noise variance, *r* is feedback noise ratio, and *g*_*c*_ is controller gain. Parameters tuned in this investigation are highlighted in red.

Motor controls are calculated by comparing the desired laryngeal state to an internal estimate of the current state of the larynx, which is maintained by a process of state prediction and subsequent correction based on auditory and somatosensory feedback signals. The controls are scaled by a tunable controller gain *g*_*c*_. An efference copy of these commands is used to predict the state of the larynx for the following time step, and consequently the expected auditory and somatosensory feedback. Meanwhile, the actual sensory feedback is generated by the larynx model. This simulated feedback can be altered to simulate the effects of a pitch perturbation experiment. Each sensory feedback signal contains Gaussian noise with zero mean and tunable variance *σ* and has some tunable delay Δ. The delayed and noisy feedback is compared with the predicted feedback to generate an error signal. The error signals from each feedback modality are used to correct the internal estimate of laryngeal state. The error signals are weighted by a Kalman gain matrix, which is calculated based on the noise in each signal, to determine the appropriate correction to the internal state estimate. Across multiple frames, the auditory output of the larynx model can be compared to a behavioral pitch perturbation response.

### State space representation

The plant of the control system, the larynx, is modeled using a state space representation of a dynamical system (Eq 1,2; [28]). Specifically, the larynx is modeled as a simple spring-mass system in which the length of the spring represents muscle tension on the vocal folds, and is therefore linearly related to vocal pitch. There exist many more complicated and accurate models of the generation of voice by the complex system of muscles controlling the larynx (see [29, 30]); however, for the current study, a very simple model was used to represent broadly the dynamics of the plant that must be controlled.

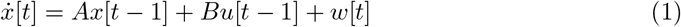

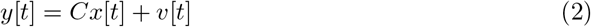

In these equations, *x* is the state (position and velocity of the mass on the spring) of the larynx dynamical system (elsewhere referred to as “larynx”), *t* is the current time step, *u* is the control input, and *w* is Gaussian process noise. The variable *y* is a vector containing auditory and somatosensory feedback and *v* is Gaussian measurement noise. *A* and *B* are matrices encoding the spring constant, damping constant, and mass of the system, which would be determined by the anatomy of the individual, and *C* encodes the transformation from state to sensory feedback. These three matrices are constants of the larynx plant (Eq 3) and remain fixed throughout the study at the values used by Houde et al. [27].

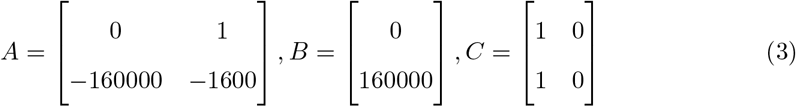

Process noise *w* is defined such that

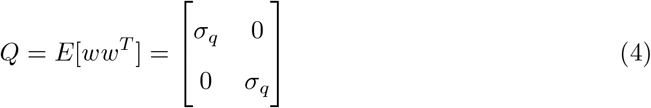

where *σ*_*q*_ is the variance of noise applied to each element of laryngeal state. *σ*_*q*_ was held constant at 1e-8 throughout the simulations, a value that was found to produce stable simulator output.

Measurement noise *v* is defined such that

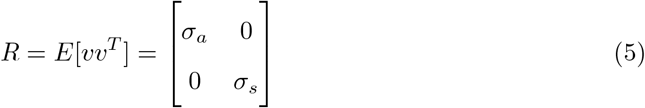

where *σ*_*a*_ is the variance of Gaussian noise on the auditory feedback signal and *σ*_*s*_ is the variance of Gaussian noise on the somatosensory signal.

Another state space system defined by *A* = 0, *B* = 1, *C* = 1 is connected in series preceding this main system, serving to integrate over the main system and thus produce a laryngeal position from the motor commands output by the main system.

Additionally, the full continuous system is discretized using zero-order hold methods with a 0.004 s sampling time.

### Pitch perturbation

The variable *y* is a two-element vector containing auditory and somatosensory feedback from the larynx. The pitch perturbation is implemented as an addend to the auditory component of the vector as follows:

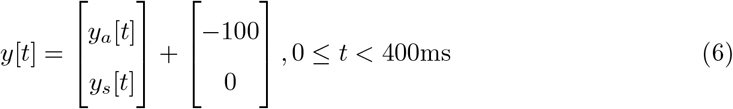

### Controller

The controller command to change the state of the larynx is defined by

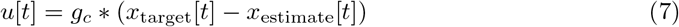

where *g*_*c*_ is the controller gain, *x*_target_ is the desired laryngeal state and *x*_estimate_ is the internal estimate of the laryngeal state.

### Observer

The observer provides the internal estimate of the state of the larynx through iterative prediction of state and subsequent update of the prediction using sensory feedback. The state is predicted by

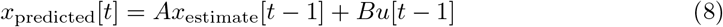

where *A* and *B* matrices are identical to those used to define the larynx (Eq 3)and *u*[*t −* 1] is the efference copy of the controller commands from the previous timestep (Eq 7). Predicted state is used to predict the sensory feedback by

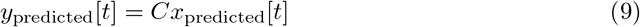

where *C* is identical to the state-to-feedback transformation matrix of the laryngeal system (Eq 3). The error between the predicted sensory feedback and the actual sensory feedback is then determined by

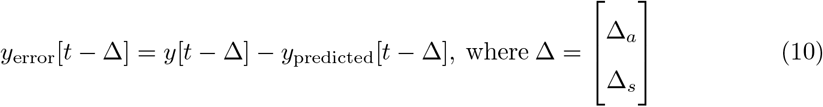

where Δ_*a*_ is auditory feedback delay and Δ_*s*_ is somatosensory feedback delay.

Sensory feedback error is used to estimate the error in laryngeal state by

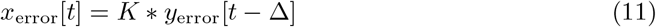

where *K* is the steady-state Kalman gain. Kalman gain is the optimal gain matrix calculated by first solving the discrete-time algebraic Ricatti equation (Eq 12) using the Python Control Systems Library (python-control; [31]) for 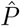 where the equation is satisfied when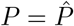.

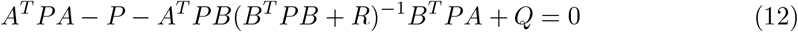

In the above equation, *A* and *B* are defined by the larynx dynamical system (Eq 3) and *Q* and *R* are defined by noise variances (Eq 4, Eq 5). Optimal Kalman gain is then calculated as follows where *C* is defined by the larynx dynamical system (Eq 3).

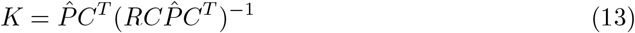

Finally, the predicted laryngeal state is updated by the error to find the estimated state using

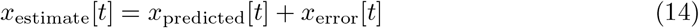

### Tunable Parameters

The values of a set of tunable parameters in the SFC model affect the shape of the model output, a simulated time course of vocal *f*_*o*_ in response to a mid-trial pitch perturbation. These parameters include the sensory delay parameters Δ_*a*_ and Δ_*s*_, controller gain *g*_*c*_, and sensory feedback noise variance parameters. In order to separate the effects of absolute noise level from the relative amount of noise between the two sensory feedback modalities, the noise variance parameters *σ*_*a*_ and *σ*_*s*_ were parameterized to *σ* and *r* such that *σ*_*a*_ = *σ* and *σ*_*s*_ = *σ/r*. Thus *σ* represents the variance of sensory feedback noise overall in both auditory and somatosensory feedback modalities and *r* represents the ratio of auditory feedback noise variance to somatosensory feedback noise variance. Overall feedback noise variance *σ* was explored on a log_10_ scale in order to more effectively search many orders of magnitude of noise variance. Thus the parameter set tuned in this investigation was *θ* = {Δ_*a*_, Δ_*s*_, *σ, r, g*_*c*_}

### Expression of Study Hypotheses in the Context of the SFC Model

Houde et al. [9] proposed two possible explanations for the behavioral pitch perturbation response differences between CA and control groups: 1) an increase in the CA group’s reliance on auditory feedback, or 2) an increase in the CA group’s reliance on all types of sensory feedback collectively compared to that of the control group.

These hypotheses can be examined using the feedback noise parameters, which are inversely related to reliance on sensory feedback through the calculation of Kalman gain *K*. An increase in the variance of the noise distribution of a particular feedback modality results in a lower Kalman gain on that feedback signal. Thus the hypotheses can be conceptualized in terms of the SFC model as follows:

*Hypothesis 1* : Compared to the control group, the CA group exhibits increased reliance on auditory feedback over somatosensory feedback. Mathematically, this can be expressed as:

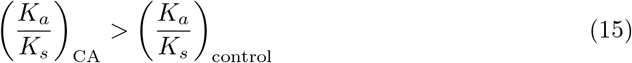

Since Kalman gain is inversely related to noise variance, this is equivalent to:

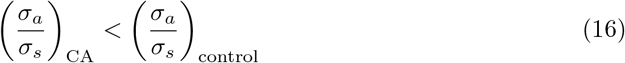

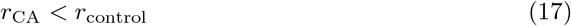

*Hypothesis 2* : Compared to the control group, the CA group displays an increased reliance on all types of sensory feedback collectively due to impairment of the feedforward system. Mathematically, this can be expressed as:

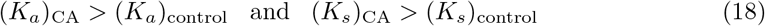

which is equivalent to:

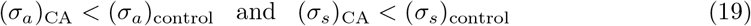

or simply:

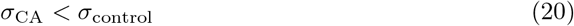

Thus Hypothesis 1 can be examined by comparing the feedback noise ratio parameter *r* between the two groups, while Hypothesis 2 can be examined by comparing the overall feedback noise variance parameter *σ* between groups. The finding of a smaller value for *r* in the CA group compared with the control group would support Hypothesis 1, while a smaller value for *σ* in the CA group would support Hypothesis 2.

## Results

### Parameter effect size

Bayesian inference was used to fit a five-parameter SFC model of voice *f*_*o*_ to the empirical pitch perturbation responses of CA and control groups as observed in Houde et al. [9]. For each empirical data set, simulation-based inference (SBI; https://github.com/sbi-dev/sbi/; [21, 22] was used to generate a posterior likelihood distribution across values for each parameter. To improve robustness, the posterior distributions from 10 repetitions of the inference procedure were combined. Fig. 3 shows the combined posterior distributions generated for each parameter, with the median and 95% Bayesian credible interval marked in each distribution. Since tests of significance lose meaning under the high statistical power of simulated data [32], effect size was used to quantify which parameters were most different between control and CA groups. Glass’s delta effect size was used because it is designed to compare populations with unequal variances. The mean and standard error of effect size calculated from 100 bootstrap samples of size 1000 are annotated on each subplot.

**Fig 3.**
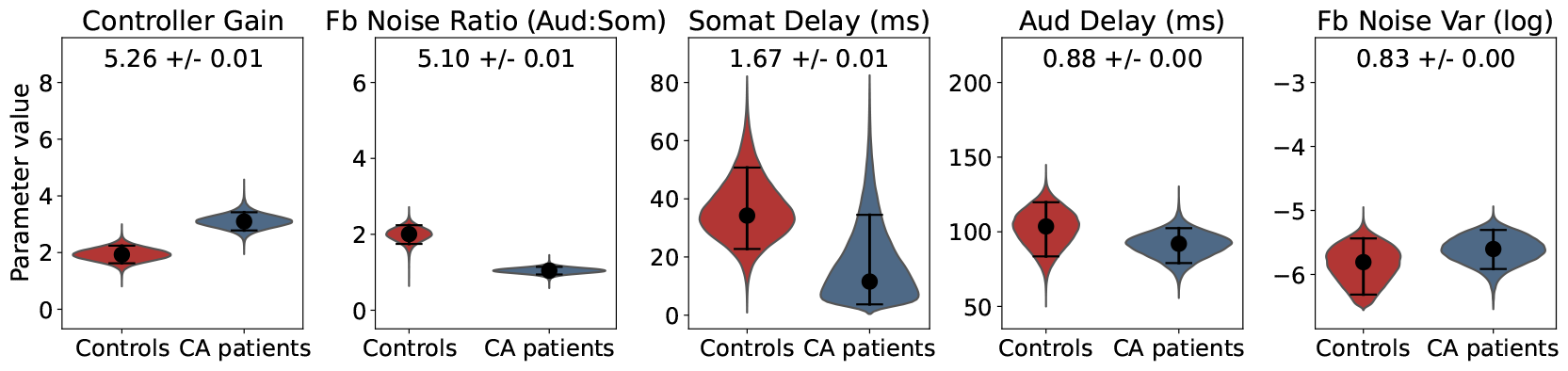
Parameter likelihood distributions. Posterior likelihood distributions (pooled from 10 repetitions of inference) for each parameter, with control group distributions on the left (red) and CA group distributions on the right (blue). The 95% Bayesian credible interval is indicated by horizontal bars and the median value of each distribution is indicated by a black dot. Glass’s delta effect size (mean and standard error from bootstrap samples) is printed at the top of each subplot.

Controller gain and feedback noise ratio had the largest effect size, while somatosensory feedback delay, auditory feedback delay and overall feedback noise variance had relatively small effect size. Notably, feedback noise ratio had a much larger effect size than the overall variance of sensory feedback noise, suggesting that the relative amount of noise between feedback modalities contributed much more to group differences than the overall level of feedback noise. All of the distributions are unimodal, which suggests there was a single optimal parameter set rather than several local optima.

### Model fit

The median of each marginal likelihood distribution was chosen as the inferred parameter set for each participant group (see Table 1). The inferred parameter sets were validated by using them in the SFC simulator to check that the results were broadly similar to the empirical data from each group. The close alignment of the tuned model with the empirical data can be seen in Fig 4. The mean of 100 simulator outputs using each group’s inferred parameter set were plotted to account for stochasticity within the SFC simulator. The standard error of these simulator outputs was also plotted, but was too small to be distinguished from the mean. Root mean square error (RMSE) between the simulator output and the behavioral data was used to quantify the quality of model fit for each group with a statistic not explicitly optimized during training. RMSE (mean ± standard error across 100 simulations) was 0.8613 *±* 0.0002 cents for the control group (4.06% of the range of the data) and 1.5544 *±* 0.0003 cents for the CA group (3.84% of the range of the data).

**Table 1.** Inferred values for control and CA groups.

| Parameter | Aud Delay (ms) | Somat Delay (ms) | Fb Noise Var ($\log_{10}$ scale) | Fb Noise Ratio (Aud:Somat) | Controller Gain |
| --- | --- | --- | --- | --- | --- |
| Symbol | $\Delta_a$ | $\Delta_s$ | $\sigma$ | $r$ | $g_c$ |
| Control | 102.7 | 35.3 | -5.8 | 2.0 | 1.9 |
| CA | 91.5 | 15.5 | -5.6 | 1.0 | 3.1 |

**Fig 4.**
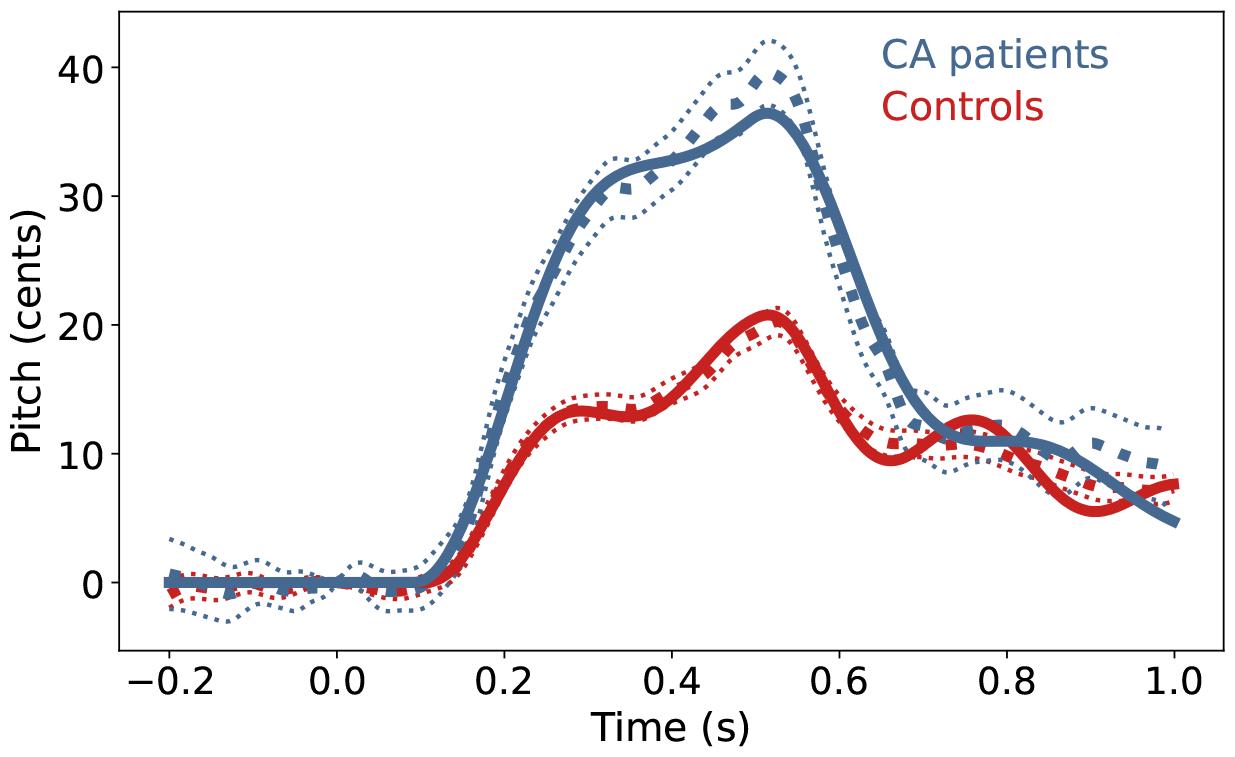
Model fit. The simulator output (solid lines) using the inferred parameter set for each group closely aligned with the behavioral data (dotted lines) previously seen in Fig 1. For simulator output and behavioral data, the mean is plotted with a thick line and standard error with a thin line, although standard error on simulator output was too small to visibly distinguish from the mean. Blue lines indicate data associated with the CA group and the corresponding simulator output, while red lines indicate those associated with the control group.

### Ablation study

Above, we have shown a successful model fit of the pitch perturbation responses for CA and control groups and quantified the parameter distribution differences between groups; however, the impact of these parameter differences on the simulator output are not immediately clear from previous results. Thus, ablation techniques [33] were used to understand the extent to which differences in inferred parameter values may translate to meaningful changes in model output. In this ablation study, the impact of each parameter on group differences was assessed by fitting the behavioral data set from the CA group with a series of reduced models. For each reduced model, one of the five original parameters was fixed to the control group’s inferred value for that parameter (as listed in Table 1) and a model composed of the four remaining parameters was used to fit the behavioral data. This allowed us to quantify the impact of each parameter on group differences and simulate the pitch perturbation response we might expect to see if individuals with CA did not differ from controls in terms of each parameter.

As seen in Fig 5, the quality of the model fit for the CA group was most greatly impacted by fixing the feedback noise ratio parameter, followed by the controller gain parameter. This provided additional evidence that these parameters were the most impactful in explaining the differences between the pitch perturbation responses of the two groups. Fixing either the somatosensory feedback delay or feedback noise variance parameter did not seem to change the quality of model fit from that of the full five-parameter model, supporting the idea that these parameters have low impact in explaining the differences between CA and control groups. Once again, the impact of the feedback noise ratio parameter was much greater than that of the feedback noise variance parameter, showing that the absolute amount of noise in each feedback signal is less impactful than the relative amount of noise between signals. To validate the method, the reduced models were also used to fit the control group data. As expected, RMSE between the simulated and behavioral control group responses remained similar to that of the full model, showing that changes in the RMSE of the CA group in the reduced model fit are attributable to group differences in parameter values rather than an artifact of reducing the number of parameters.

**Fig 5.**
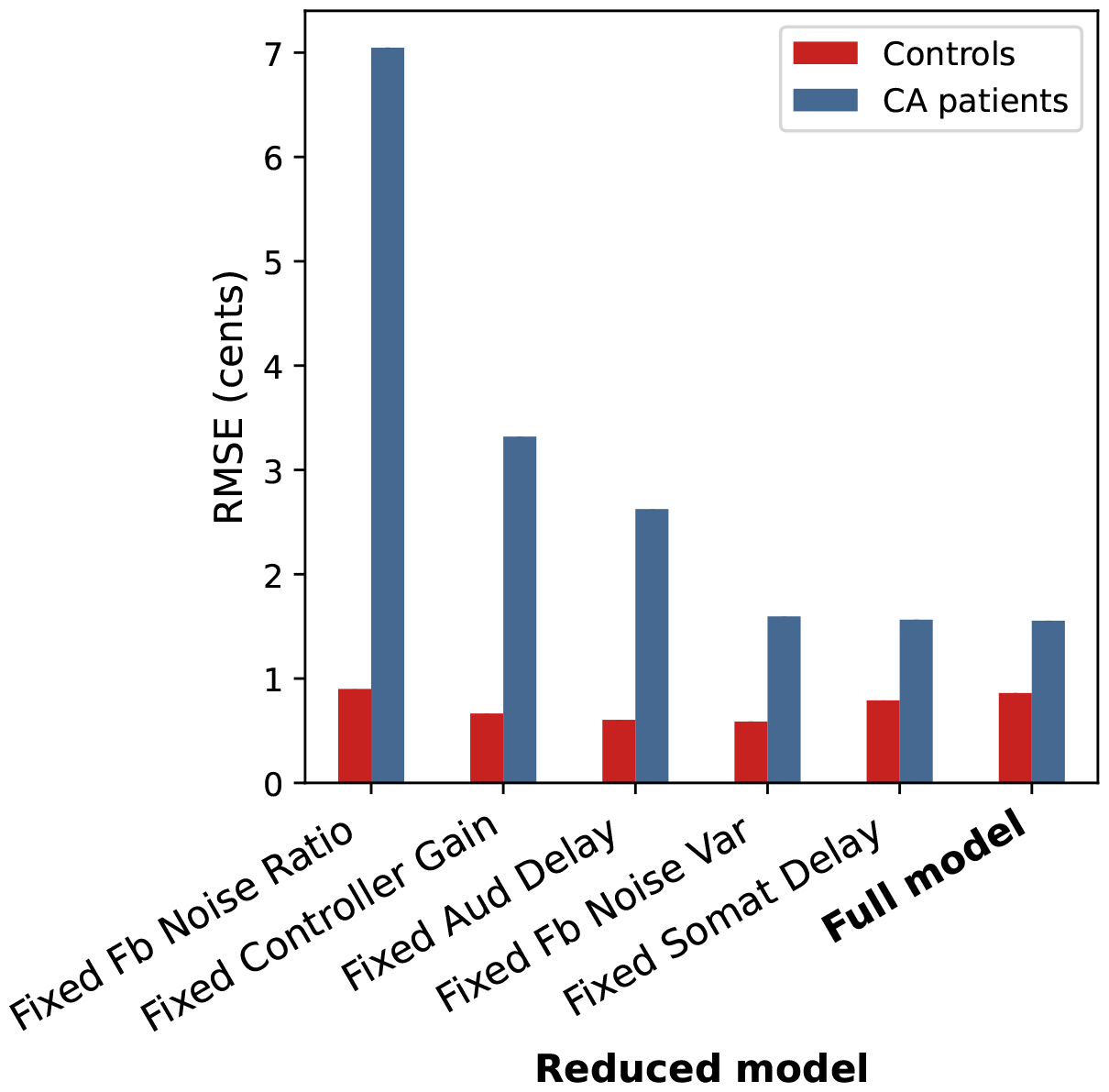
Quality of fit for reduced models. Quality of fit for each 4-parameter model with one parameter fixed to the inferred value for the control group. The quality of each model’s fit is quantified by RMSE between empirical data and optimized model output (mean RMSE across 100 simulator runs).

For each four-parameter model, the optimized simulator output of control and CA group data are shown in Fig 6, along with the posterior distributions of the remaining parameters. The optimized simulator output for each four-parameter model provides insight regarding the impact of each parameter on the shape of the pitch perturbation response, while the distributions of the remaining parameters indicate which parameters may interact with the ablated parameter.

**Fig 6.**
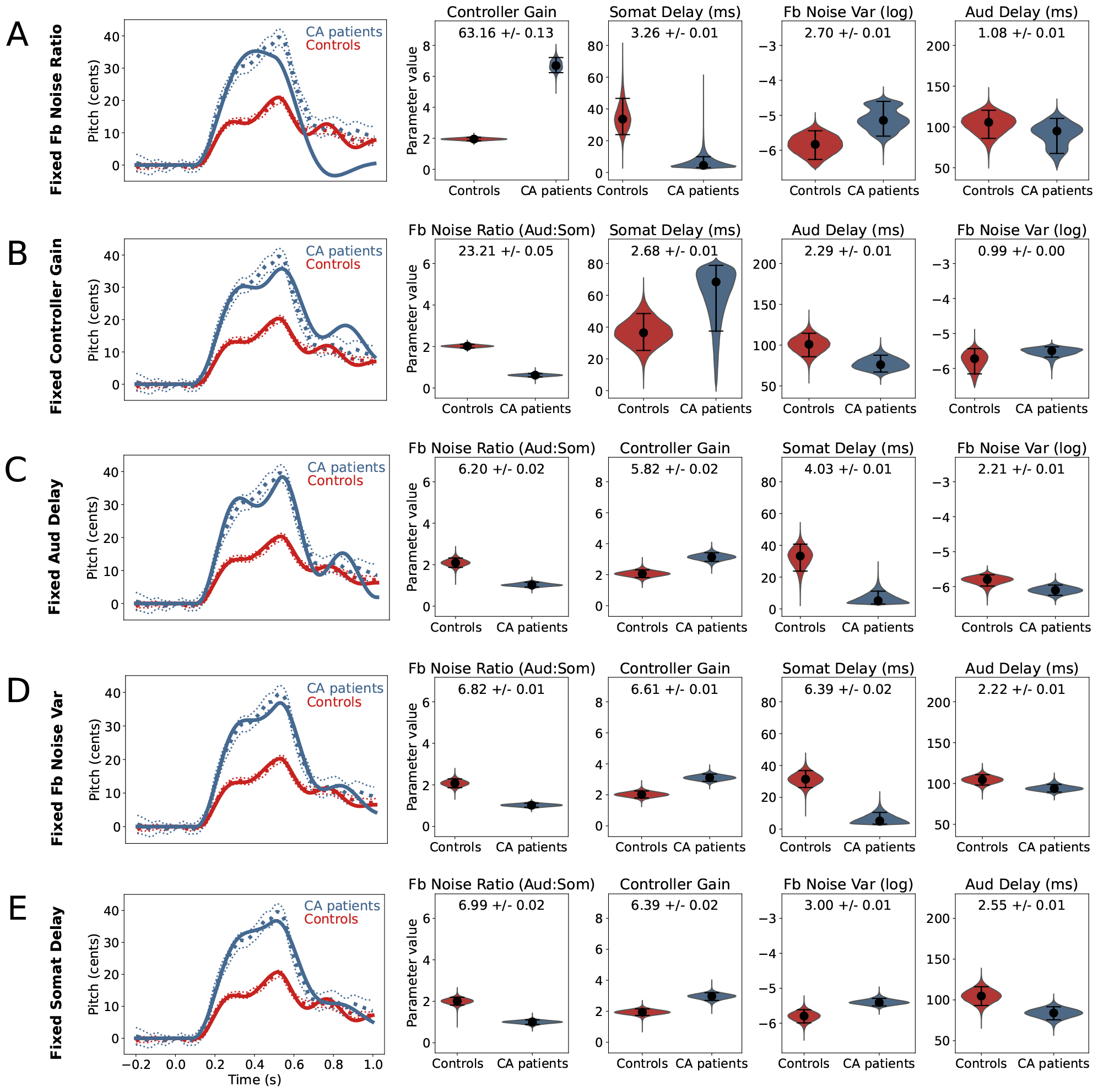
Reduced models: simulator output and posterior distributions of remaining parameters. Inference results for each of five reduced models in which a single parameter is fixed to the inferred value of the control group from the full model fit and the remaining four parameters are used to fit the behavioral pitch perturbation data. Shown for each reduced model are the posterior distributions of the four remaining parameters and the simulator output (mean of 100 simulations) using the inferred reduced parameter set.

The results of the ablation of the feedback noise ratio parameter are shown in Fig 6A. We can see from the posterior distributions that while the somatosensory feedback delay, feedback noise variance, and auditory feedback delay parameter distributions remained mostly similar to those of the full model, the controller gain parameter distribution for the CA group shifted up to a much larger median value. This allowed the simulated response to nearly reach the peak response of the behavioral CA data, but it dipped far below the behavioral CA data during the period from 0.7 to 1.0 s after perturbation onset.

Alternately, when the controller gain parameter was ablated, the feedback noise ratio greatly increased in effect size as the median value of the CA group distribution shifted down from 1.04 to 0.62. The median value of the somatosensory delay parameter also increased but the effect size of this parameter increased only slightly due to the large variance of the distribution. These changes allowed the simulator output to nearly reach the peak magnitude of the behavioral data of the CA group, but with the controller gain fixed at the value of the control group, the optimized simulator output was higher and more oscillatory than the behavioral data during the period from 0.7 to 1.0 s after perturbation onset. Thus contributions from both the feedback noise ratio and controller gain parameters are needed to replicate both the large magnitude of the CA group response and the flat, moderate time course of *f*_*o*_ after the end of the perturbation.

When auditory feedback delay was fixed to the inferred value for the control group (Fig 6C), the CA group response became more oscillatory, contributing to a small increase in error from the behavioral data. Fixing either somatosensory feedback delay or overall feedback noise variance, meanwhile, did not appear to change the model fit. It can be seen in (Fig 6D) that the ablation of feedback noise variance caused a change in the distribution of the somatosensory feedback delay parameter, and vice versa in (Fig 6E). Thus these two parameters, interestingly, seem to have similar effects on model output.

## Discussion

In this study, simulation-based Bayesian inference was used to disambiguate possible mechanisms underlying the observed differences in the pitch perturbation responses of individuals with CA and healthy controls by identifying speech motor control model parameter differences between these two groups. The results indicate that most of the differences between the pitch perturbation responses of individuals with CA and healthy controls can be explained by differences in the feedback noise ratio and controller gain. These parameters showed both (a) the largest effect size between groups in comparing the posterior distributions and (b) the greatest loss of fit accuracy for the CA group when fixed to the inferred value of the control group.

Our finding that the feedback noise ratio was lower in the CA group than in the control group with substantial effect size supports Hypothesis 1, which suggests increased reliance on auditory feedback relative to somatosensory feedback in the CA group. In contrast, we did not find evidence in support of Hypothesis 2, the idea that CA group displays an increased reliance on all types of sensory feedback collectively. The small effect size for the overall sensory feedback noise parameter and the minimal change when this parameter was ablated were inconsistent with this hypothesis.

Here we will discuss how these findings fit into the previous literature regarding pitch perturbation response in individuals with CA, new hypotheses generated by the model, and the strengths and limitations of the modeling techniques used in this analysis.

### Evidence for overreliance on auditory feedback

The high effect size of the feedback noise ratio suggests that the CA group may show increased reliance on auditory feedback relative to somatosensory feedback. This may appear to conflict with the results of Li et al. [10], which showed in individuals with spinocerebellar ataxia a decreased cortical P2 response in the right superior temporal gyrus (STG), primary auditory cortex (A1), and supramarginal gyrus (SMG) during a pitch perturbation task. However, while this finding may provide evidence against the idea that the absolute value of auditory feedback error gain is larger in the CA group than the control group ((*K*_*a*_)_CA_ *>* (*K*_*a*_)_control_), we argue that it does not rule out the idea of increased sensitivity to auditory feedback as we have defined it in this study, that is, relative to somatosensory feedback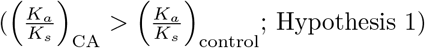. If both auditory and somatosensory gains are smaller in the CA group compared to controls, it is possible that the resulting ratio between auditory gain and somatosensory gain may still be larger in the CA group compared to the control group (i.e., if the somatosensory gain in the CA group were much smaller compared that of the control group). Defining feedback noise in terms of a ratio between sensory modalities has been previously used by Crevecoeur et al. [23] to model relative sensitivity to visual and proprioceptive feedback in arm reaching. Furthermore, the P2 response may not be a direct measure of gain on auditory feedback error; the functional relevance of P2 response remains controversial and has been suggested to have contributions from multiple sensory modalities including auditory and somatosensory [34]. Although this view is highly speculative, we argue that the measurement of decreased P2 response in the CA group [10] does not completely rule out Hypothesis 1 and that our findings warrant further investigations.

### Lack of evidence for overall overreliance on feedback

Given the ties between internal models and the cerebellum [19, 20], it is indeed somewhat surprising that we did not find evidence of overreliance on sensory feedback overall. It would be reasonable to expect that if the internal models, and thus the state prediction process, are disrupted in CA, sensory feedback would be more heavily weighted to compensate for impairment in the feedforward system [9]. However, the results indicate similar weighting of sensory feedback overall between the two groups, with perhaps even slightly lower weighting (increased noise) of sensory feedback in the CA group. We speculate that perhaps the nature of the disruption of internal models impairs the integration of sensory feedback, which has similar effects to a slightly decreased weighting. Alternatively, perhaps the auditory nature of the task created a result in which auditory feedback appears to be emphasized rather than overall feedback.

### Novel hypothesis generated by the model: controller gain parameter

In addition to support for the previously stated hypothesis that individuals with CA show increased reliance on auditory feedback relative to somatosensory feedback, our results lead us to propose an additional hypothesis to be tested in future investigations. The high impact of the controller gain parameter suggests a difference between individuals with CA and healthy controls in the scaling of the motor command to the larynx, which has likely neural substrate in the motor cortex [25]. Although any direct ties between this parameter and the cerebellum are unknown, we speculate that perhaps the increased value of controller gain in the CA group, which could also be interpreted as increased sensitivity to target error (the difference between the intended production and the actual production) may indicate a learned mechanism in the motor cortex to compensate for deficits related to the disorder. Further study is needed to investigate this hypothesis.

### Efficacy of simulation-based inference for model parameter estimation

Simulation-based inference allowed us to obtain posterior likelihood distributions across each parameter rather than a single set of optimal values. This made it possible to take into account the spread of each distribution in addition to the inferred value and thus determine the effect size of the difference between groups for each parameter.

Additionally, in a complex system it is possible to have many locally optimal parameter sets that can achieve high quality fit to the empirical data. Obtaining a unimodal posterior likelihood distribution for each parameter showed that the parameter set selected could approximate a globally optimal solution within the bounds of the prior. Simulation-based inference also offered advantages over other Bayesian methods since it does not require an analytical form of the model (allowing us to analyze a complex model like state feedback control) and is more computationally efficient than other numerical techniques such as Markov Chain Monte Carlo methods [35].

### Limitations

It is important to note that the current work is beneficial for generating hypotheses, rather than drawing conclusions. The larynx was modeled here as a simple spring-mass system. While this implementation may approximate many of the dynamics of laryngeal movement, future studies should investigate how results may vary with more detailed models of laryngeal muscles. Additionally, real speakers vary in their vocal tract dimensions, damping, and other properties that are not possible to measure. The model cannot quantitatively reflect all of these properties to represent an actual speaker.

Furthermore, these properties were left unchanged in the model throughout the present set of simulations, so the vocal tract properties of at most one speaker were represented. Meanwhile, the behavioral data used in the present study were group averages of many subjects with different physiologies. To account for the averaging of multiple observations, the simulator was run 100 times and the mean output was plotted.

However, standard error on model output was so small (*<* 0.1 cent) that it could not be differentiated from the mean production on the plot. Standard error on the empirical mean was larger, in the range of 0-3 cents, further demonstrating that the model does not account for the variation that is present among individual speakers.

Additionally, the present study was limited to the model changes that could be captured by five model parameters. The same model architecture was used to model both control and CA groups, with all differences captured in the tuning of model parameters. Thus possible differences in the structure of neural systems between the groups were not tested in the current study. Furthermore, many of the model parameters tested here are abstract concepts whose precise neural implementation is not yet fully understood. Each parameter that is represented in the model as a single value may in fact be the result of many complex processes. Finally, only five parameters were considered in the present study. Since training data was generated by testing values for each parameter in combination with all other parameters, the number of parameters tested was in exponential tradeoff with the resolution of values tested and the range of the prior for each one. While the irrelevance of other untested model parameters cannot be proven, the tested parameters can be argued to be sufficient since the model output for each set of inferred values closely matched the behavioral data.

### Conclusion

This work has shown that the controller gain and feedback noise ratio parameters have high effect size and large contributions to the group differences between the pitch perturbation responses of individuals with CA and healthy controls. These results (a) provide support for the previous hypothesis that individuals with CA show increased sensitivity to auditory feedback prediction error and (b) generate a new hypothesis of increased sensitivity to target error in the CA group. Furthermore, this work demonstrates the value of simulation-based inference methods in analyzing behavioral speech data using tunable parameters of a state feedback control model.

## Materials and Methods

### Simulation-based inference overview

The SBI package in Python (https://github.com/sbi-dev/sbi/; [21, 22]) was used to obtain posterior likelihood distributions of five tunable parameters in the SFC model (see Model section for an overview of SFC) for the CA and control group average pitch perturbation responses observed in Houde et al. [9]. As detailed in Fig 7, SBI takes as input a computational model with a finite set of input parameters (“simulator”), a prior distribution for each parameter, and an empirical observation analogous to the output of the simulator. It generates a data set by running the simulator with inputs from the prior distributions of the parameters, and then, using the Sequential Neural Posterior Estimation (SNPE) option for inference, trains a deep neural density estimator to predict the posterior distribution of parameters given the empirical observation.

**Fig 7.**
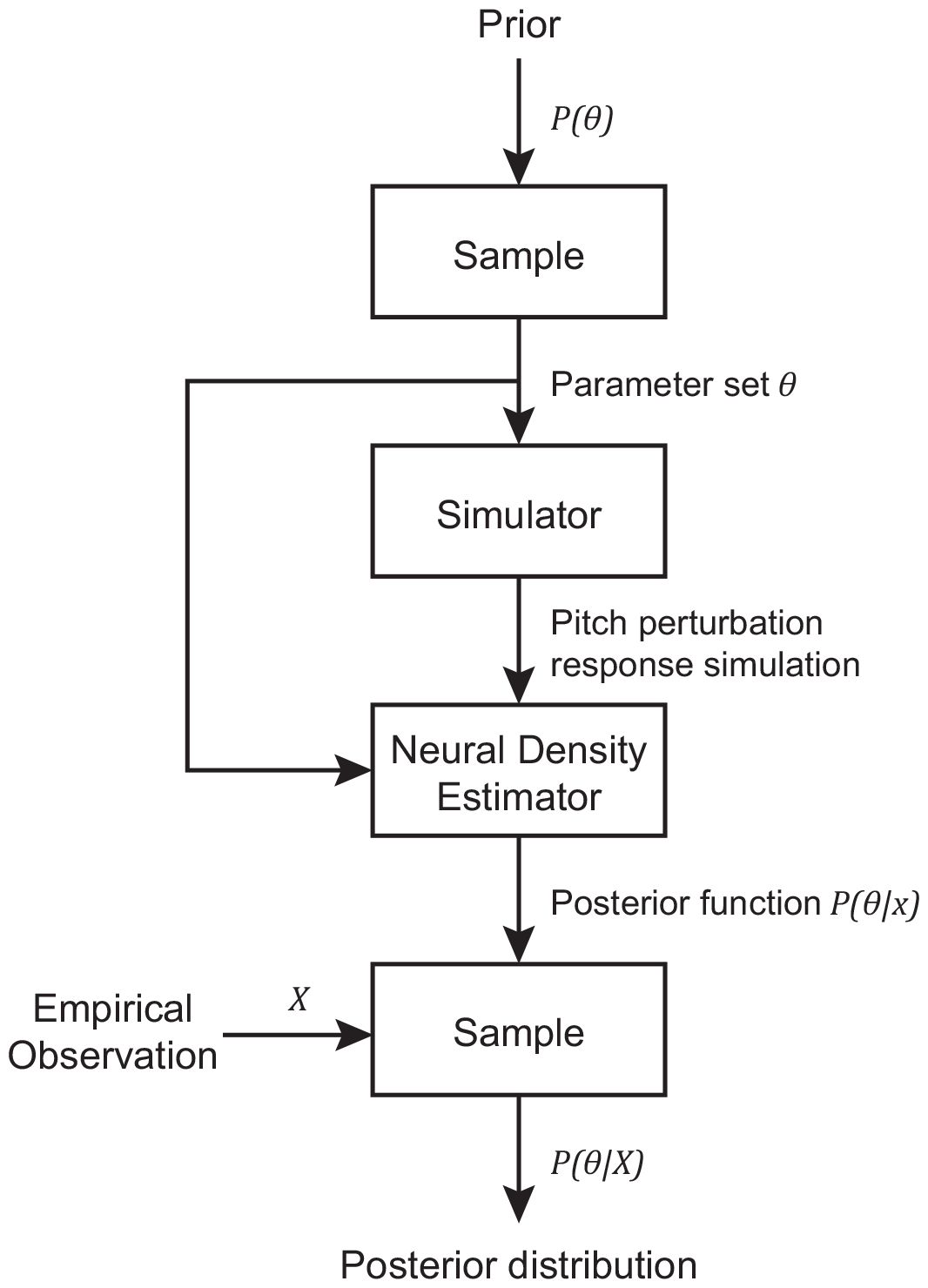
SBI overview. A pipeline for performing simulation-based inference [21, 22]. Parameter values are inferred for a particular empirical observation given a mechanistic model (“simulator”) and a prior distribution for each tunable parameter.

### Simulator

A Python implementation (https://github.com/jessicagaines/1d-sfc) of a state feedback control model of *f*_*o*_ [25, 27] *was used as the simulator (see Model section for more details). The input parameter set included the following parameters: auditory feedback delay Δ*_*a*_, somatosensory feedback delay Δ_*s*_, controller gain *g*_*c*_, feedback noise variance *σ*, and feedback noise ratio *r*. The parameters *σ* and *r* are parameterizations of auditory feedback noise variance *σ*_*a*_ and somatosensory feedback noise variance *σ*_*s*_ such that *σ*_*a*_ = *σ* and *σ*_*s*_ = *σ/r*. This idea of exploring relative levels of feedback noise between different sensory modalities was also used by Crevecoeur et al. [23] in their state feedback control model of arm reaching. The output simulated a time course of voice *f*_*o*_ in response to a -100 cent, 400 ms, mid-utterance perturbation of *f*_*o*_ feedback. Random noise was added to model output during training as Jin et al. [36] found that this increased reconstruction accuracy. Uniform noise distributions of increasing width were tested and since the quality of model fit stopped improving for training noise distribution wider than 7 cents, noise with distribution *∼ U* (*−*3.5, 3.5) was added to the simulator output during training.

### Prior distribution of parameters

The SBI inference procedure requires the input of a prior distribution for each parameter, which defines the search space of the training data. A uniform prior was used for each parameter. For some parameters, the bounds of the prior could be estimated from measurable quantities. For example, Abbs and Gracco [37] indicate that latencies in response to a somatosensory perturbation are on the order of tens of milliseconds, so a prior of 3 to 80 ms was used for somatosensory delay. Latencies to auditory perturbations, meanwhile, have been reported in the range of 100-200 ms [38], so a prior of 50 to 200 ms was used. Measurable latencies may be greater than delays since they include motor response time, so the lower bounds were set lower than the measured response latencies. The lower bound for somatosensory delay is quite low, in the range of what is typically associated with non-cortical reflex [39], but since no minimum delay value can be definitively measured, we opted not to restrict the prior based on this information. The remaining parameters could not be estimated from measurable quantities, so wide initial priors were selected to fully explore the space. Initial bounds of 0.1 to 10 were selected for the feedback noise ratio and controller gain parameters to include two orders of magnitude, and the range of 1e-10 to 1e-1 was selected for feedback noise variance. The auditory feedback noise variance parameter was converted to a base-10 logarithmic scale to search many orders of magnitude more effectively. For likely parameter sets in this regime, the simulator was found to be unstable for feedback noise variance less than 1e-6.5, and so the lower bound for this parameter was increased to 1e-6.5 to train the likelihood estimator on stable simulator outputs. Finally, the results of the wide prior showed that the tails of the posterior likelihood distributions of feedback noise variance, feedback noise ratio, and controller gain parameters were far from the upper bounds of each prior. The bounds were narrowed slightly to increase the search resolution for each parameter and decrease the computational resources needed. The final bounds selected are shown in Table 2. This choice of prior is validated by the result that the likelihood distributions for each parameter (see Fig 3) lie comfortably within these bounds, except for feedback noise variance, which was restricted for stability, and somatosensory delay, which by definition cannot be less than one frame of simulator operation.

**Table 2.** Uniform priors with the following bounds were selected for each parameter.

| Parameter | Aud Delay<br>(ms) | Somat Delay<br>(ms) | Fb Noise Var<br>(log<sub>10</sub> scale) | Fb Noise Ratio<br>(Aud:Somat) | Controller<br>Gain |
| --- | --- | --- | --- | --- | --- |
| Symbol | $\Delta_a$ | $\Delta_s$ | $\sigma$ | $r$ | $g_c$ |
| Lower<br>bound | 50 | 3 | -6.5 | 0.1 | 0.1 |
| Upper<br>bound | 200 | 80 | -3 | 6 | 8 |

### Empirical observation

Behavioral group average pitch perturbation responses from CA and control groups [9] as described in the Introduction were used as empirical observations to sample the posterior. Each behavioral data set was downsampled from 413 to 300 frames per 1.2 s trial to match the output of the simulator.

### Inference

10^5^ simulations were used to train the neural density estimator [36]. The posterior was then sampled 10^4^ times for each group. To improve robustness, this procedure was repeated 10 times and the samples from each repetition were pooled to obtain the final parameter distributions. A 95% Bayesian credible interval was calculated for each distribution. Glass’s delta was used to calculate the effect size of each parameter between groups. To assess the quality of model fit, the median of each pooled distribution was considered the ”inferred value” for each parameter and each inferred parameter set was supplied as input to the simulator. To reduce the effects of stochasticity within the simulator, 100 simulations were run with each inferred parameter set and the mean of these was plotted. The quality of the fit was assessed quantitatively by calculating the point-wise root mean square error (RMSE) between the model output and the empirical data. This statistic was not used in training the neural network and therefore provided a separate method of quantifying the success of the model fit.

### Ablation study

Finally, an ablation study was used to further understand the impact of each parameter on model output [33]. One at a time, each parameter was ablated by fixing it to the inferred value (the median value of the pooled posterior distribution) of the control group and repeating the inference procedure to generate posterior distributions for the four remaining parameters. The medians of these distributions were once again used in the simulator to assess the quality of fit for each reduced model using RMSE and compare the result to that of the full model. A greater increase in error for a particular reduced model indicated that the parameter ablated in that model had greater impact on group differences.

